# A hole in Turing’s theory: pattern formation on the sphere with a hole

**DOI:** 10.1101/2022.11.10.515940

**Authors:** Johannes G. Borgqvist, Philip Gerlee, Carl Lundholm

**Affiliations:** Wolfson Centre for Mathematical Biology, Mathematical Institute, University of Oxford, Andrew Wiles Building Radcliffe Observatory Quarter (550) Woodstock Road, Oxford, OX2 6GG, Oxfordshire, United Kingdom; Mathematical Sciences, University of Gothenburg, Chalmers tvärgata 3, Gothenburg, SE-412 96, Västra Götaland, Sweden; Mathematical Sciences, Chalmers University of Technology, Chalmers tvärgata 3, Gothenburg, SE-412 96, Västra Götaland, Sweden; Department of Mathematics and Mathematical Statistics, Umeå University, MIT Building, 3rd floor Linneaus Väg, Umeå, SE-907 36, Västerbotten, Sweden

**Author notes:** Contributing authors.

**Keywords:** Turing patterns, RD-models, Bud scars, FEM

## Abstract

The formation of buds on the cell membrane of budding yeast cells is thought to be driven by reactions and diffusion involving the protein Cdc42. These processes can be described by a coupled system of partial differential equations known as the Schnakenberg system. The Schnakenberg system is known to exhibit diffusion-driven pattern formation, thus providing a mechanism for bud formation. However, it is not known how the accumulation of bud scars on the cell membrane affect the ability of the Schnakenberg system to form patterns. We have approached this problem by modelling a bud scar on the cell membrane with a hole on the sphere. We have studied how the spectrum of the Laplace–Beltrami operator, which determines the resulting pattern, is affected by the size of the hole, and by numerically solving the Schnakenberg system on a sphere with a hole using the finite element method. Both theoretical predictions and numerical solutions show that pattern formation is robust to the introduction of a bud scar of considerable size, which lends credence to the hypothesis that bud formation is driven by diffusion-driven instability.

## 1 Introduction

A fascinating class of biological phenomena accounting for the emergence of complex patterns is that of *reaction diffusion* (RD) processes. Using two simple principles corresponding to chemical reactions between proteins and their movement due to diffusion, complex patterns in the concentration profile of these proteins emerge for certain types of reactions, and this pattern formation is often described by a class of *partial differential equations* (PDEs) which we will refer to as RD-models. The theoretical basis for this phenomenon, that was initially proposed by Alan M. Turing [1], is called *diffusion driven instability* [2], and based on this phenomenon RD-models have been applied in numerous situations including that of patterns in animal coatings and among fish [3], to name but a few.

Diffusion-driven instability has also been suggested to operate on the intracellular scale, in the context of cell division in the baker’s yeast *Saccharomyces cerevisiae*, also referred to as *budding yeast*. This type of yeast divides through a process called budding, where a smaller daughter cells grows out of the larger mother cell, and the spatial location on the cell membrane where the daughter cell grows out is determined by a protein called Cdc42. More precisely, in the G1-phase of the cell cycle, the activated form of Cdc42 diffuses on the cell membrane and ultimately accumulates at a specific spot called a *pole*. It is at the pole that budding occurs and the daughter cell grows out from the mother cell during the cell division. After cell division the pole turns into a bud scar, a circular region into which proteins cannot move [4].

The process of pole formation is thought to be described by an RD-process confined to the membrane of the cell. There are numerous RD-models of Cdc42-mediated cell polarisation [5–7] and these are all of activator-inhibitor type. This means that Cdc42 is shuffled between its active and inactive forms through chemical reactions, and these reactions in combination with the diffusion of the two forms of Cdc42 results in the formation of a pole due to diffusion driven instability. The simplest canonical activator-inhibitor RD-model which can form patterns is called the Schnakenberg model [8], and this model is frequently used for analysing pattern formation caused by diffusion driven instability both analytically and numerically.

One of the challenges when studying diffusion driven instability for activator-inhibitor RD-models is the gap between the theoretical predictions and numerical simulations. In general, the theoretical formulae based on linear stability analysis for diffusion driven instability are limited to relatively low-dimensional as well as geometrically simple domains, whereas simulations typically can account for a higher geometrical complexity. For example, it is not uncommon that the linear stability analysis is conducted in one spatial dimension, e.g. on a line, and then these theoretical results guide RD-simulations on a geometrical approximation of the cell in three spatial dimensions. This is problematic since there is a difference in geometry between the analytical and numerical spatial domains, both in terms of dimensionality and curvature.

The simplest non-trivial spatial domain approximating the surface of a cell is the sphere, where theoretical predictions in combination with simulations have been conducted using the Schnakenberg model by Chaplain et al. [9]. In that context, it was possible to derive theoretical thresholds for a critical rate of diffusion as well as a critical reaction strength parameter which could be used to predict the properties of the resulting pattern.

From a mathematical point of view, we can model such a bud scar on the cell membrane by a hole on the sphere together with *Neumann boundary conditions*, i.e., zero-flux. Previous theoretical results as well as previously applied numerical methods (e.g. spectral methods of lines as used in [9]) are unable to predict pattern formation in this case. With this in mind we set out to combine theoretical results from spectral theory with a finite element method (FEM) for solving the Schnakenberg system on the sphere with a hole, in order to describe the impact of bud scars on bud formation.

In this work, we have theoretically and numerically analysed the effect of a hole on the unit sphere *S*^2^ on the pattern formation of the Schnakenberg model. The emergence of patterns is controlled by the eigenvalues of the Laplace– Beltrami operator on the sphere with a hole, and we have made use of results from spectral theory to calculate approximate values of the eigenvalues as a function of the radius of the hole. The theoretical results show that the patterns that appear on the sphere are retained although a hole of considerable size is introduced. This observation was verified by solving the Schnakenberg system numerically using the finite difference scheme 1-SBEM [10] in time together with the finite element method in space. Our results show that diffusion driven bud formation appears to be robust to the presence of bud scars.

## 2 Results

### 2.1 Validation of the numerical method

To study the pattern formation on a sphere we consider the Schnakenberg model. In its dimensionless form, this model is given by the following RDsystem:

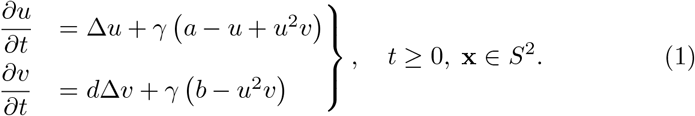

Here, *a, b, γ, d >* 0 are positive constants, *u*(**x, t**) is the concentration profile of the activator, *v*(**x, t**) is the concentration profile of the inhibitor and Δ is the Laplace-Beltrami operator on the sphere. To numerically solve (1) and thus simulate pattern formation, we implemented a numerical method that uses the finite difference scheme 1-SBEM [10] in time together with the finite element method in space. To validate the numerical method and its implementation (see the Materials and Methods section for details), we replicated the results of Chaplain et al. (see figure 4.3 in [9]) where the dynamics of the Schnakenberg model (1) on the sphere was solved numerically. Specifically, in these simulations, the following parameters were used:

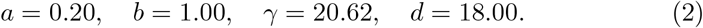

These values were chosen so that *n* = 2 is the only mode in the unstable interval (See Materials and methods for the definition of the unstable interval and how the values of *d* and *γ* were chosen). For these parameter values, the steady-states are given by

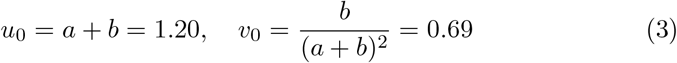

and starting from these initial conditions Chaplain et al. ran the simulations until a dimensionless time of *t* = 50 was reached. Using the same exact parameter values, we validated our implementation by reproducing the results in [9] (see Fig. 1 and Fig. 4.3 in [9]).

**Fig. 1.**
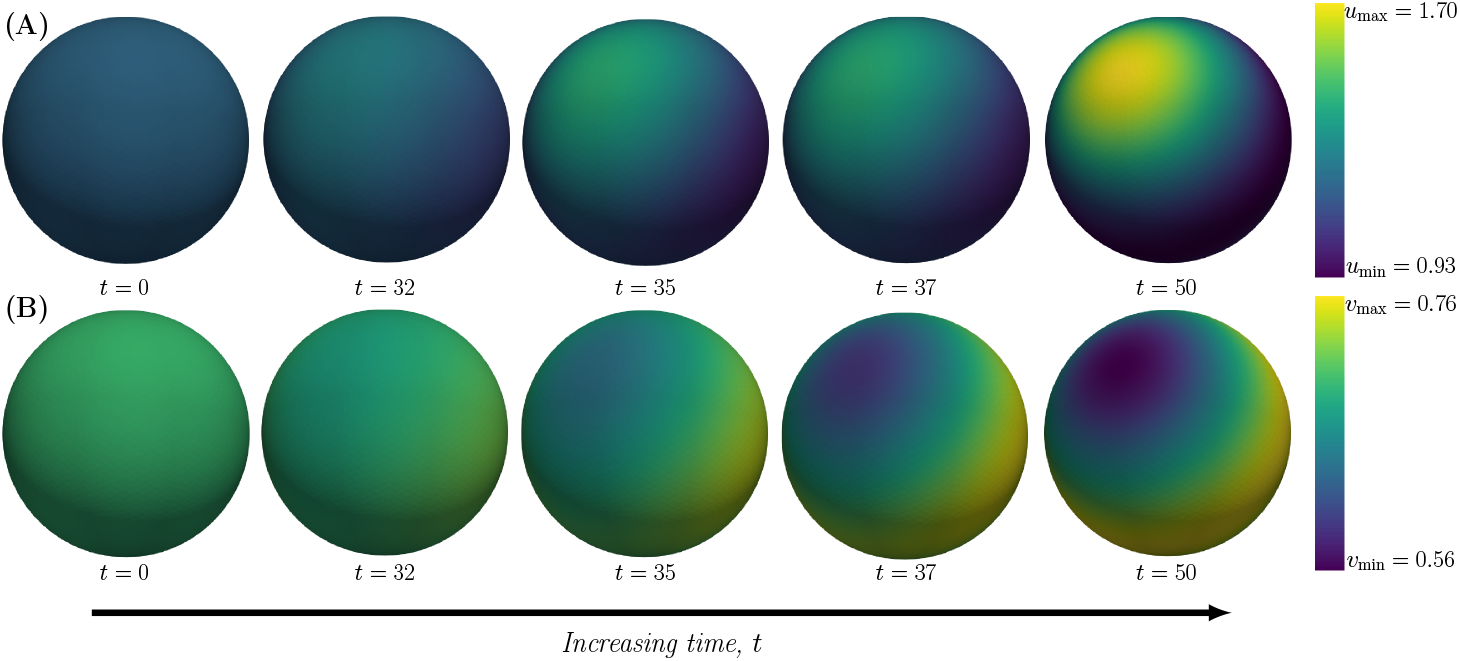
Validation of the implementation of the numerical method. The Schnakenberg model is simulated on the unit sphere *S*^2^ with the following parameter values: (*a, b, d, γ*) = (0.20, 1.00, 18.00, 20.62). The time is increasing from left to right, and the concentration profiles at the time points *t* = 0, *t* = 32, *t* = 35, *t* = 37 and *t* = 50 are shown in two cases. (**A**) The active component *u*(**x**, *t*) with a minimum concentration of *u*_min_ = 0.93 and a maximum concentration of *u*_max_ = 1.70. (**B**) The inactive component *v*(**x**, *t*) with a minimum concentration of *v*_min_ = 0.56 and a maximum concentration of *v*_max_ = 0.76. The time scale and the concentration ranges of the two species agree with Fig. 4.3 in [9].

Given the validation of our numerical implementation, we now proceed to the problem of pattern formation on the sphere with a hole.

### 2.2 Robustness of Turing patterns on the sphere with a hole: predictions from spectral theory

We model the addition of a bud scar on the cell membrane by considering the unit sphere *S*^2^ with a hole. More precisely we consider a spherical cap Ω_*ε*_ centred at the North Pole (0, 0, 1) ∈ ℝ^3^ of geodesic radius *π* −*ε*. Here *ε >* 0 correspond to the geodesic radius of the hole, and we denote the boundary of the hole by *∂*Ω_*ε*_.

It is known that as budding yeast cells go through multiple cell division the bud scars tend to aggregate on the cell membrane. Here we only model the addition of a single bud scar, but given their vicinity in real cells [4] we model the accumulation of bud scars by considering a single hole of increasing radius *ε*.

Since the formation of Turing patterns is determined by the spectrum of the Laplace–Beltrami operator (see Methods for details), we are interested in the following eigenvalue problem

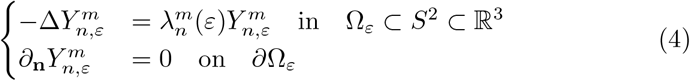

where 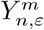 are the eigenfunctions of the Laplace–Beltrami operator on Ω_*ε*_, 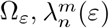 the corresponding eigenvalues, and *∂*_**n**_ is the derivative in the direction of the outer normal. Here we consider Neumann (i.e. no-flux) boundary conditions meant to describe a situation where no proteins can enter the bud scar.

Given this problem formulation, we are now interested in the eigenvalues 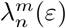 in (4) as a function of the radius of the hole *ε*. Asymptotic expansion of these eigenvalues in the limit of *ε* →0 have been derived by Bandle et al. [11]. These perturbed eigenvalues are given by [11]:

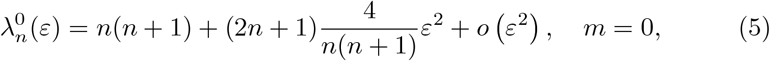

and

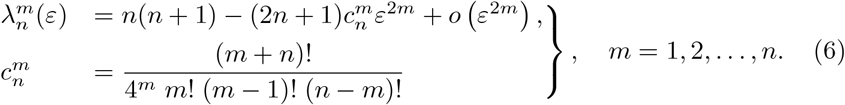

It is worth pointing out that the spectrum is continuous with respect to the hole radius and that 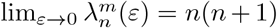, which equals the eigenvalues of the Laplace–Beltrami operator on the sphere.

In the limit of a small hole the formation of Turing patterns can thus be determined by considering a modified version of the unstable range of eigenvalues:

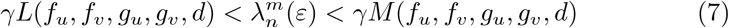

where only eigenvalues/functions that fall in the above interval contribute to spatial patterning.

Fig. 2 shows 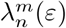 as a function of *ε* for *n* = 1 and *n* = 2, where lower and upper bounds in (7) are shown as dashed lines (see Methods for details on parameter values). For *ε* = 0 the eigenvalues for each *n* are degenerate, but as the radius of the hole increases they diverge, but remain within the pattern formation range for small values of *ε*. Thus we conclude that for these parameter settings, where a single eigenvalue lies in the unstable interval, we expect pattern formation to be unaffected by the introduction of a small hole.

**Fig. 2.**
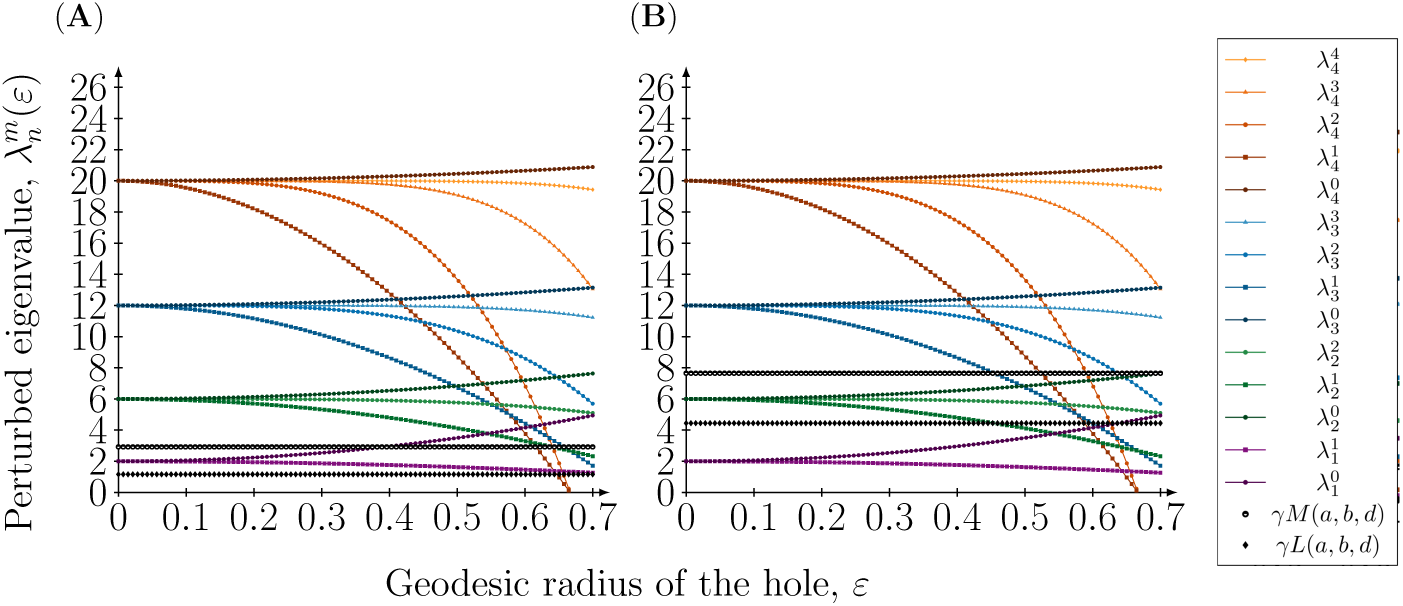
Perturbed eigenvalues as a function of the geodesic radius of the hole on the sphere. The perturbed eigenvalues 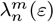 are plotted against the hole radius *ε* when *n* = 1, 2, 3, 4 and *m* = 0, 1, …, *n*. Also, the upper boundary *γM* (*a, b, d*) and the lower boundary *γL*(*a, b, d*) in the Turing condition involving the eigenvalues giving rise to patterns are illustrated in the dashed lines. The parameters defining these boundaries are chosen to (*a, b*) = (0.20, 1.00) and the value of *γ* is set to the critical value, i.e., *γ* = *γ*_*c*_(*n*) for a particular eigenmode *n*. The upper boundary *γM* (*a, b, d*) and the lower boundary *γL*(*a, b, d*) are illustrated in two cases: (**A**) (*n, γ, d*) = (1, 6.87, 20.00) and (**B**) (*n, γ, d*) = (2, 20.62, 18.00).

We now move on to investigate this theoretical prediction using a numerical implementation of the Schnakenberg model on the sphere with a hole.

### 2.3 Numerical solutions verify the theoretical prediction

To investigate the effect of a hole on pattern formation, and to test the above derived theoretical predictions, we solved the Schnakenberg model numerically on spheres with an increasing hole radius for four parameter sets, corresponding to patterns with a single excited mode (*n* = 1, 2, 3 and 4). The rate parameters *a* and *b* were fixed to the values *a* = 0.20 and *b* = 1.00 in all simulations according to (2). We set the value of the reaction strength *γ* and the relative diffusion *d* according to critical values such that a single mode was excited (see Methods for details).

An example of the effect of increasing the hole radius is shown in Fig. 3, where the mode *n* = 2 is excited and *ε* = 0 corresponds to a complete sphere without a hole. From this it appears as if the addition of a hole has little effect on the resulting pattern.

**Fig. 3.**
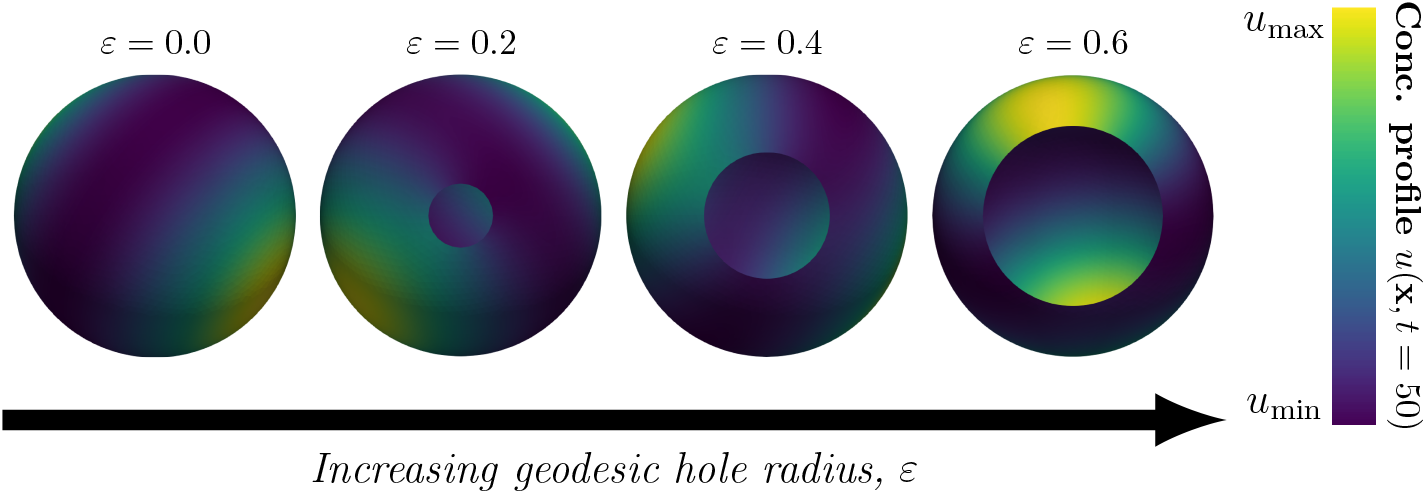
The pattern is robust with respect to the introduction of a hole in the domain. The concentration profile of the active component at time *t* = 50, denoted by *u*(**x**, *t* = 50), **x** ∈ Ω_*ε*_, is illustrated on four different meshes with a single hole located at the South Pole with geodesic radii *ε* = 0.00, 0.20, 0.40, 0.60. The parameters of the Schnakenberg model which were used to generate the above results were (*a, b, d, γ*) = (0.20, 1.00, 18.00, 20.62), and in all cases the initial conditions were set to a small perturbation around the steady-state concentrations of the two species in each node of the mesh. The maximum and minimum concentrations for the different geodesic hole radii *ε* from left to right are given by: (*ε, u*_min_, *u*_max_) = (0.00, 0.93, 1.75), (*ε, u*_min_, *u*_max_) = (0.20, 0.91, 1.74), (*ε, u*_min_, *u*_max_) = (0.40, 0.90, 1.73) and (*ε, u*_min_, *u*_max_) = (0.60, 0.93, 1.6).

In order to investigate this further, we varied the hole size determined by the geodesic radius *ε* in the range *ε*∈ [0, 0.7], and to account for the stochasticity of the solutions, which are introduced via the initial conditions, each simulation was repeated 20 times. We set out to characterise the resulting patterns both by projecting the solutions onto the spherical harmonics and by quantifying the number of poles, the maximum concentration of *u*, the relative area of the poles and the minimum distance from the hole to a pole (see Methods for details).

The projection of the numerical solution onto the spherical harmonics provides a way of investigating which modes are excited in the resulting pattern. Fig. 4 shows the decomposition in the case where *n* = 1 is excited. Here we observe large variability in the coefficients 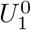 and 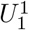, whereas the other coefficients show less variability and take values close to zero. An exception is the coefficient of the constant mode 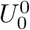, but this value corresponds to the steadystate value from which the pattern emerges, and can be explicitly calculated in the following way: the zeroth mode is stable with respect to spatial perturbations and we therefore expect it to remain approximately constant as the dynamics evolve. Since it is constant we can express it in terms of the initial concentration 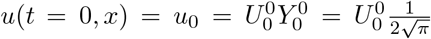, and since *u*_0_ = 1.2 we obtain 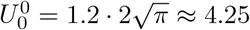, which is close to the value observed for small *ε* in Fig. 4A.

**Fig. 4.**
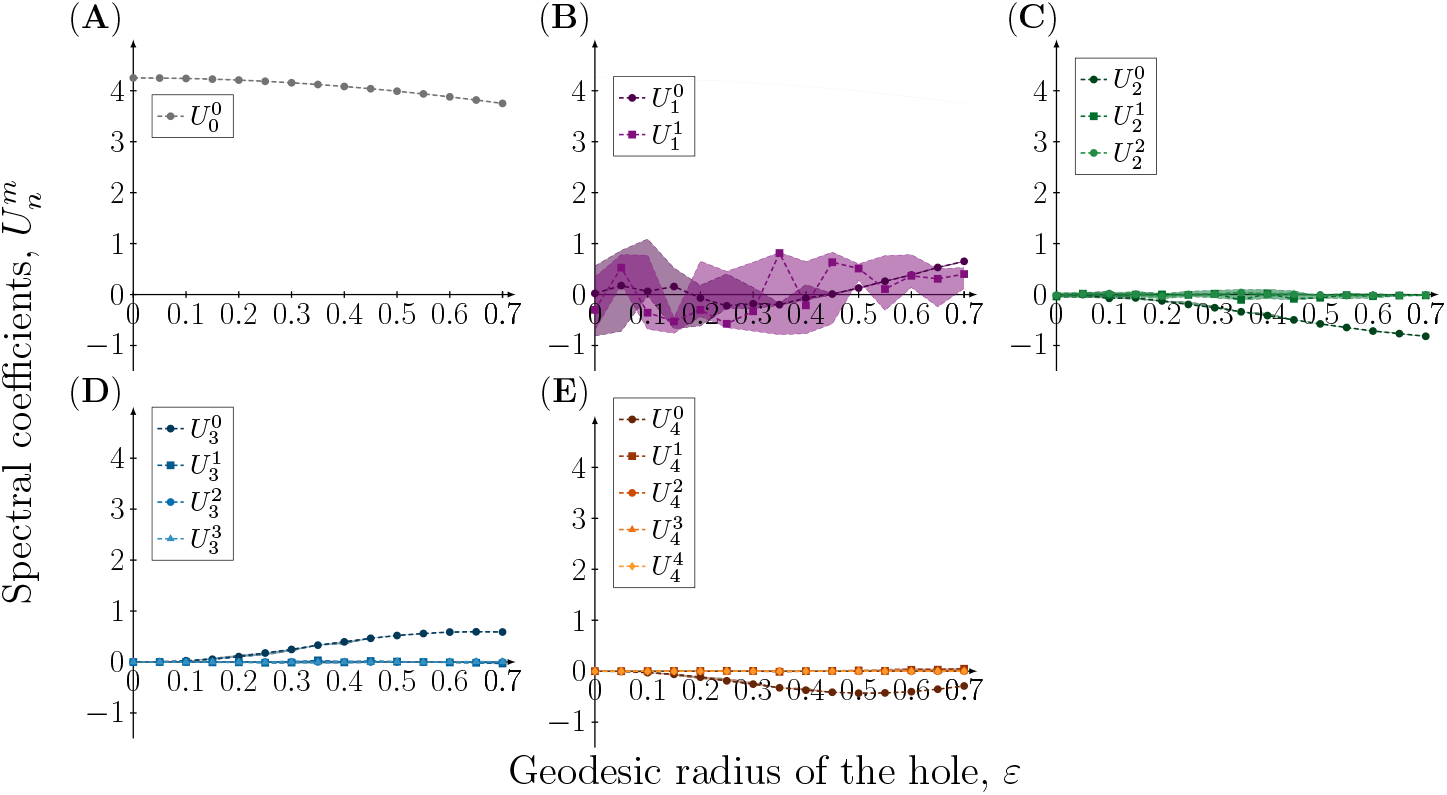
Spectral coefficients of the concentration profile of the active component at time t = 50 on meshes with a single hole with increasing radius when (n, d) = (1, 20). The coefficients of the eigenfunctions 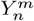 in the spectral approximation of the concentration profile *u*(**x**, *t* = 50), **x** ∈ Ω_*ε*_ resulting from the rate parameters (*a, b, d, γ*) = (0.20, 1.00, 20.00, 6.87) are plotted as a function of the geodesic radius of the hole *ε*. Due to the stochasticity in the initial conditions, each simulation has been repeated 20 times, and to account for the variation in the coefficients the 95%, 50% and 5% percentiles are plotted.

The case where *n* = 2 is excited for the sphere is shown in Fig. 5. Here we observe a similar pattern with variation in the coefficients corresponding to the excited mode and all other coefficients remaining small, with the exception of 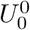. The pattern is repeated for *n* = 3 and 4 (see Supplementary material), and we thus conclude that variation occurs only in the mode that was excited for the sphere and that introducing a hole has a minor effect on other modes. We now turn to our quantitative metrics of pattern formation. Fig. 6A shows that for *n* = 1 the number of poles that are present at *t* = 50 in the numerical solution is constant and equal to one for the entire range of hole sizes. Also the relative pole area (Fig. 6B) and the maximum concentration of *u* (Fig. 6C) are preserved. In contrast, the distance from the centre of the hole to the closest pole varies considerably. Precisely the same pattern is observed for *n* = 2 (see Fig. 7) (similar results for *n* = 3 and 4 can be found in the Supplementary material).

**Fig. 5.**
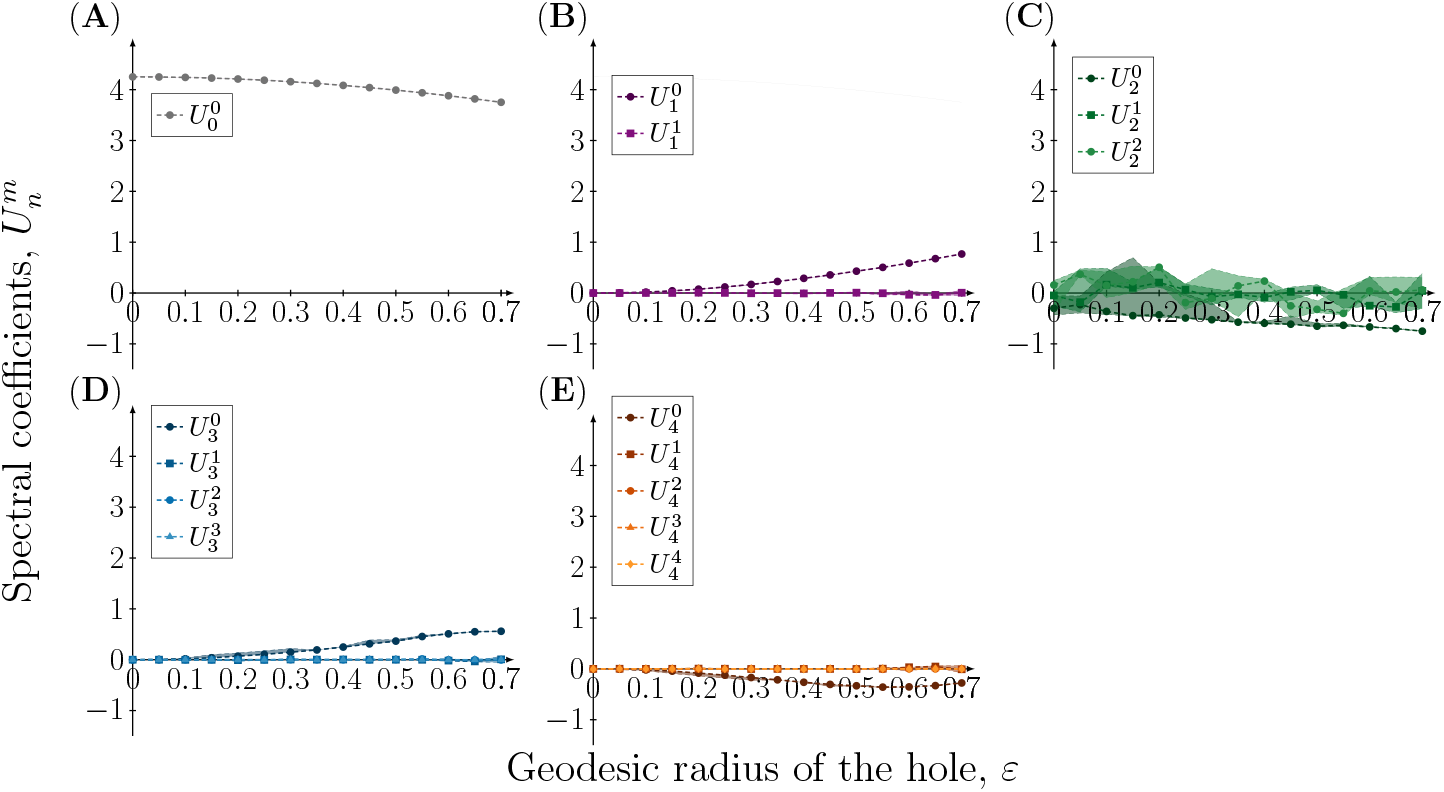
Spectral coefficients of the concentration profile of the active component at time t = 50 on meshes with a single hole with increasing radius when (n, d) = (2, 18). The coefficients of the eigenfunctions 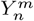 in the spectral approximation of the concentration profile *u*(**x**, *t* = 50), **x** ∈Ω_*ε*_ resulting from the rate parameters (*a, b, d, γ*) = (0.20, 1.00, 18.00, 20.62) are plotted as a function of the geodesic radius of the hole *ε*. Due to the stochasticity in the initial conditions, each simulation has been repeated 20 times, and to account for the variation in the coefficients the 95%, 50% and 5% percentiles are plotted.

**Fig. 6.**
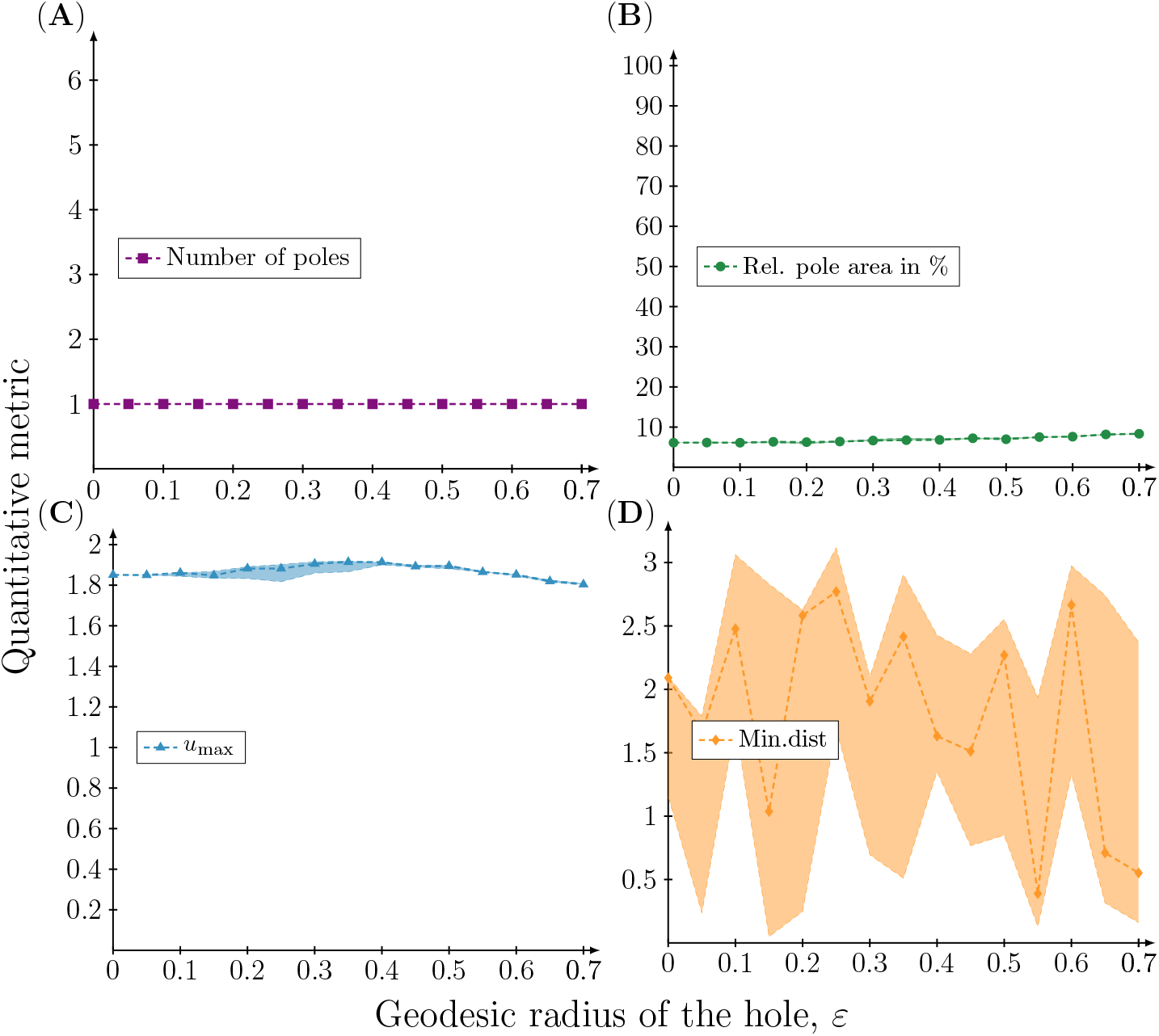
Quantitive metrics of the concentration profile of the active component at time t = 50 on meshes with a single hole with increasing radius when (n, d) = (1, 20). Four different quantitative metrics of the concentration profile *u*(**x**, *t* = 50), **x** ∈Ω_*ε*_ resulting from the rate parameters (*a, b, d, γ*) = (0.20, 1.00, 20.00, 6.87) are plotted as a function of the geodesic radius of the hole *ε*. (**A**) The number of poles corresponding to high concentration regions. (**B**) The total pole area relative to the total surface area. (**C**) The maximum concentration *u*_max_. (**D**) The minimal *great circle distance* between a pole and the hole. Each simulation has been repeated 20 times and therefore the 95%, 50% and 5% percentiles of the quantitative metrics are plotted.

**Fig. 7.**
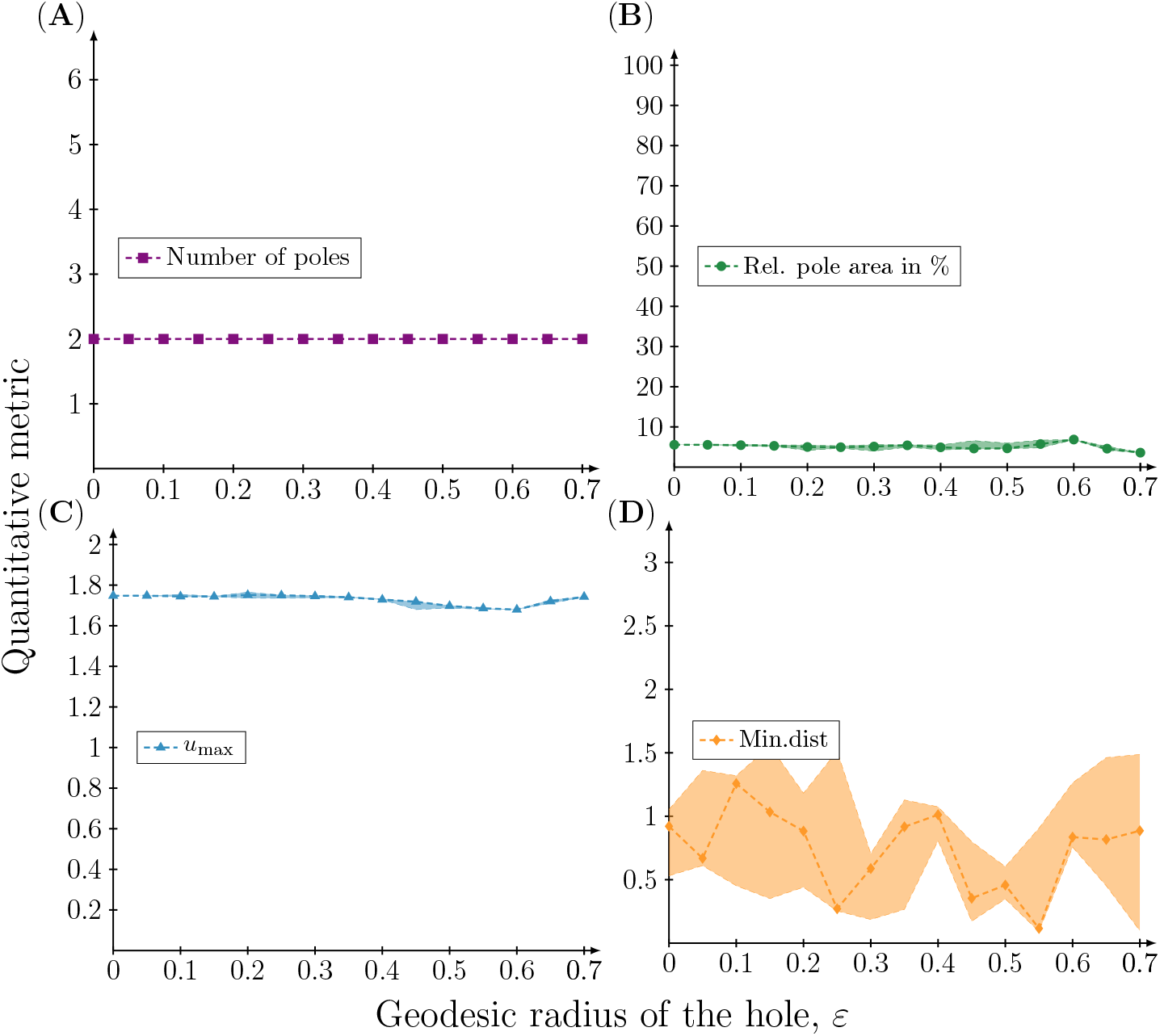
Quantitive metrics of the concentration profile of the active component at time t = 50 on meshes with a single hole with increasing radius when (n, d) = (2, 18). Four different quantitative metrics of the concentration profile *u*(**x**, *t* = 50), **x** ∈ Ω_*ε*_ resulting from the rate parameters (*a, b, d, γ*) = (0.20, 1.00, 18.00, 20.62) are plotted as a function of the geodesic radius of the hole *ε*. (**A**) The number of poles corresponding to high concentration regions. (**B**) The total pole area relative to the total surface area. (**C**) The maximum concentration *u*_max_. (**D**) The minimal *great circle distance* between a pole and the hole. Each simulation has been repeated 20 times and therefore the 95%, 50% and 5% percentiles of the quantitative metrics are plotted.

These results show that the introduction of a hole does not affect the resulting pattern. The variation of the distance between the hole and closest pole appears simply because the orientation of the pattern depends on the random initial conditions and thus varies between simulations. This also explains the variation observed in the spectral coefficients of the excited mode. The orientation of the pattern causes the coefficients of the excited modes that constitute the pattern to vary, when in fact the pattern (modulo rotation) is preserved.

## 3 Discussion

In this work, we have investigated the effect of introducing a hole of varying size on the unit sphere on the Turing patterns exhibited by the Schnakenberg model. Using a FEM-based implementation together with a spectral analysis in terms of the excited eigenfunctions of the Laplace–Beltrami operator contributing to the solution, we concluded that the quantitive properties of the final patterns are largely conserved when a hole is introduced.

This observation can be in part explained by considering an asymptotic expansion of the eigenvalues of the Laplace–Beltrami operator on the sphere with a hole. In the limit of small hole sizes, the perturbed eigenvalues can be written as a power series in the hole radius *ε*. This implies that the eigenvalues are continuous with respect to the hole radius and that the eigenvalues that are in the unstable range for the sphere (without a hole), remain so even in the presence of small hole (see Fig. 2).

The opposite also holds true, for small *ε* no other eigenvalues enter the unstable region, and thus the pattern is conserved. The conclusion hinges on the continuity of the spectrum of the Laplace– Beltrami operator with respect to the removal of a small circular region of the sphere. Despite the fact that the topology of the domain is altered, the spectrum is continuous and 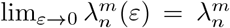. This property holds for a much wider class of perturbations of the domain, as has been proven by Chavel & Feldman [19], and in particular the addition of finitely many holes on the sphere. However, explicit formulae for the perturbed eigenvalues have only, in the more general setting, been derived for Dirichlet boundary conditions. In the reaction diffusion setting, considered here, it makes less sense to consider such boundary conditions, yet we conjecture that pattern formation in the Schnakenberg model (with Neumann boundary conditions) will remain intact even in the case of finitely many holes of small radius.

There are are a number limitations to our approach and the conclusions drawn from it. In order to analyse the resulting pattern we project the numerical solutions on the sphere with a hole onto the spherical harmonics, although they do not form an orthonormal ℒ_2_ basis for the domain. However, we expect this to be a reasonable approximation, in particular for small hole sizes. This implies that the spectral decomposition shown in Fig. 4 and Fig. 5 should be interpreted with caution, especially for hole sizes in the upper range.

The asymptotic expansions obtained from Bandle et al. [11] are only valid in the limit of small hole sizes. This means that our conclusions regarding which eigenvalues that fall within the unstable region (7) are only strictly valid in the limit of small hole sizes. Further, it implies that the plots showing the relation between perturbed eigenvalues and the unstable region (Fig. 2) are not accurate for large hole sizes.

Motivated by the fact that bud scars of ageing yeast cells tend to cluster [4], we have considered a hole of increasing radius meant to represent several smaller bud scars. This is a crude approximation, but the analytical results mentioned above suggest that the effect of several small holes on the eigenvalues of the Laplace–Beltrami operator should have a similar impact as single hole.

Our results suggest that bud formation in budding yeast is robust to changes that alter the topology of the membrane on which the reactions that drive the accumulation of Cdc42 occur. Bud formation corresponds to an instability of the eigenvalue *n* = 1 whose eigenfunction contains a single peak, but we have shown that patterns corresponding to *n* = 1, 2, 3 and 4 are stable with respect to the addition of a hole as well.

From a biological stand point this is far from surprising since budding yeast cells are known to form many buds (and thus contain many bud scars) before they reach senescence and stop dividing. From a mathematical modelling point of view our results lend credence to the idea that activator-inhibitor dynamics (as modelled by the Schnakenberg model) are a good model of the process of bud formation. If the dynamics were highly sensitive to the addition of a hole they would indeed present a poor model of the phenomenon.

In relation to this it should be noted that budding yeast cells of advanced age are dysfunctional and tend to form multiple buds [20–22]. This can in part be explained by the fact that cell size increases with age, and in the non-dimensional Schnakenberg model this corresponds to an increase in the reaction strength *γ*. Mathematically, this leads to a widening of the unstable region in the spectrum, which might cause additional eigenmodes to become excited leading to patterns with multiple poles. Additionally, it has been hypothesised that oxidative stress alters the enzymatic activity of the proteins that move Cdc42 between its active and inactive form. In terms of the Schnakenberg model this would correspond to changing the reaction parameters *a* and *b*, which could disrupt the pattern forming properties of the system. However, as we have shown, the addition of a hole perturbs the eigenvalues, which might contribute to additional eigenmodes being excited. Thus, it might be both the increasing cell size, altered enzymatic activity and a large number of bud scars that contribute to dysfunctional budding.

A feature not captured by our model is that new buds tend to form close to existing bud scars [21]. This could be the results of space dependent reaction rates [23], which could be introduced into our model. Further, it would be of interest to investigate if other RD-models such as the Thomas model [24] and the Gray-Scott model [25] are equally robust to the addition of holes on the sphere.

In conclusion, we have shown that RD-dynamics for pole formation in budding yeast can be modelled with a finite element approach, which makes it possible to account for the presence of an existing bud scar of varying size. This method is highly flexible and can be adapted to solve RD-models on any domain that can be represented by a mesh. Numerical solutions show that the introduction of a bud scar does not affect the resulting pattern, and this can in part be explained by appealing to results from spectral theory, that describe how the eigenvalues of the Laplace–Beltrami operator are perturbed in the presence of a hole. The stability of pole formation with respect to the addition of bud scars add theoretical evidence for pole formation being a Turing pattern of activator-inhibitor type, but more research in this direction is needed to explain dysfunctional pole formation in ageing cells and also the clustering of bud scars on the cell membrane.

## 4 Methods

All the scripts for generating the results presented in this work can be accessed through the public GitHub repository associated with this work [12]. The entire project is written in Python and it only uses open-source dependencies. To visualise these results and to generate the figures, the software ParaView [13, 14] has been used.

### 4.1 Turing patterns on the sphere

It has been hypothesised that the formation of buds in budding yeast, which initiates the formation of a daughter cell and subsequent cell division, is driven by reaction and diffusion of molecules that are confined to the cell membrane. The spatial distribution of chemical species on the membrane can be described by a coupled set of partial differential equations known as reaction-diffusion equations. The formation of spatial patterns in such systems is known as a Turing instability and can appear under specific constraints on the diffusion coefficients and reaction rates of the involved chemical species.

A reaction-diffusion system of two interacting species *u* and *v* on the sphere *S*^2^ can be written in non-dimensonal form as

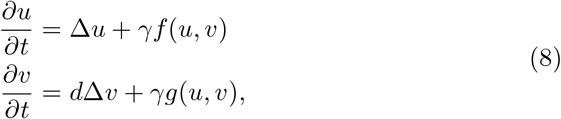

where *d* is the ratio of the (dimensional) diffusion coefficients, *f, g* the reaction rates, *γ* is a constant related to the radius and Δ is the Laplace–Beltrami operator on *S*^2^.

A homogeneous steady state (*u*_0_, *v*_0_) exhibits diffusion-driven Turing instability [2, 9] if it satisfies *f* (*u*_0_, *v*_0_) = *g*(*u*_0_, *v*_0_) = 0 and the following inequalities are satisfied

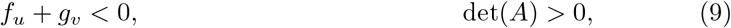

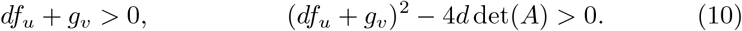

Here *f*_*u*_, *g*_*v*_ are partial derivatives evaluated at (*u*_0_, *v*_0_), and *A* is the Jacobian, also evaluated at the steady state. The type of pattern that appears depends on the range of unstable modes in the spectrum of the Laplace–Beltrami operator. The Laplace–Beltrami eigenvalue problem on the sphere is:

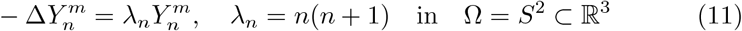

Here, the eigenfunctions are the spherical harmonics 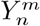, where |*m*| ≤ *n*. A mode *n* is unstable if it falls in the range [2, 9]

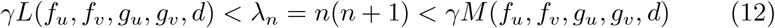

where

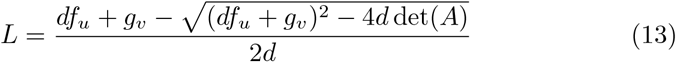

and

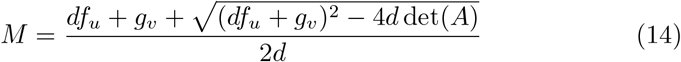

If there exists at least one such *n* then the homogeneous steady state is unstable with respect to spatial perturbations and we can expect a heterogeneous pattern composed of the eigenfunctions 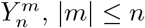, that satisfy (12).

In summary, provided that the parameters in the reaction terms *f, g* of the RD-model (8) are chosen so that the Turing conditions (9) and (10) are satisfied, it is the spectrum of the Laplace–Beltrami operator that determines pattern formation. More specifically the eigenmodes *n* that satisfy the bounds in (12).

### 4.2 Finding critical values for *γ* and *d*

In order to obtain diffusion-driven instability the diffusion coefficient *d* needs to be above a certain critical value, which is given by [9]:

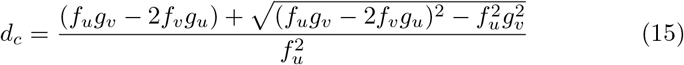

where *f*_*u*_, *f*_*v*_, *g*_*u*_, *g*_*v*_ correspond to the partial derivatives in the Jacobian evaluated at the steady state. Only the eigenvalues that lie within the unstable interval contribute to pattern formation, and in order to isolate a single eigenvalue (and thus a single mode) one sets *γ* to a critical value which is a function of the eigenvalue *n* [9]:

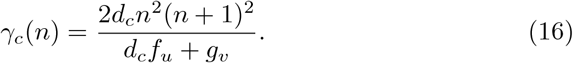

For the parameters *a* = 0.20 and *b* = 1.00, the critical diffusion is given by *d*_*c*_ ≈17.01. If we pick *γ* = *γ*_*c*_(*n*) for a specific eigenmode *n* and *d* sufficiently close to *d*, then the resulting concentration profile will be a linear combination of the eigenfunctions corresponding to the particular eigenmode *n* (in addition to the zeroth eigenmode) and no other eigenmodes. To this end, we have picked *d* sufficiently close to *d*_*c*_ as well as the value *γ* = *γ*_*c*_(*n*) in the four cases corresponding to *n* = 1, 2, 3, 4, respectively, in our experiments.

### 4.3 The mesh for geometrically approximating the sphere with a hole

The meshes were generated using Gmsh [15]. We generated our holes in the mesh by intersecting a cylinder with the sphere, and then we removed the intersection in order to obtain a mesh of the spherical cap.

Given these meshes, we implemented our FEM-based numerical scheme for solving the Schnakenberg model on a spherical mesh with a single hole located at the South Pole, i.e., centered at (*x, y, z*) = (0, 0, −1). A visualisation of the meshes can be found in the Supplementary information (Fig. S1).

### 4.4 The numerical scheme for simulating Schnakenberg’s RD-model using a FEM-FD approach

We implemented the 1-SBEM numerical scheme [10] for solving the Schnakenberg model in FEniCS [16, 17]. For all details of this algorithm, we refer to [10], but here follows a short summary of the involved steps.

Given the Schnakenberg RD-model, the first step is to multiply the system of PDEs with two test-functions 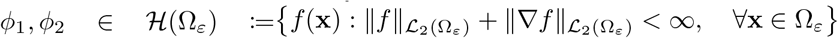 where the ℒ_2_-norm is induced by the inner product

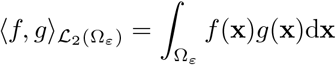

and then integrate over the domain Ω_*ε*_. Given this operation, the RD-system is written as follows:

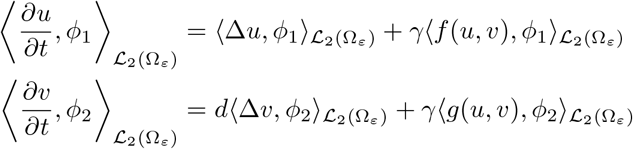

where the reaction terms *f, g* are given in Equation (1). Using Green’s formula on the diffusive terms in combination with the fact that we have a Neumann boundary condition on the boundary of the hole *∂*Ω, we can move all terms to the left hand sides and then add the two equations. This results in one single equation which is our *variational formulation*:

Find *u*(·, *t*), *v*(·, *t*) ∈ ℋ (Ω_*ε*_) for a fixed *t* ∈ ℝ such that

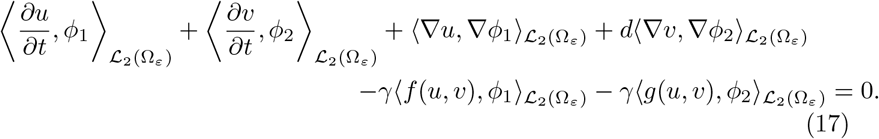

To find the numerical approximation, this variational formulation is restricted to a discrete subspace of continuous piecewise linear functions. This method is referred to as *continuous Galerkin of order 1* denoted by cG(1).

To approximate the time derivatives in Equation (17), we use a finite difference scheme. Given that we want to solve the original RD system for times in the time interval [0, *T*] for some final time *T >* 0, we define a partition τ_*k*_ of the interval [0, *T*] into *N* pieces for some integer *N >* 0 as follows:

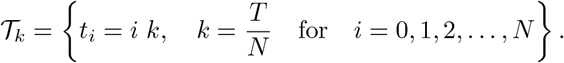

Given this partition, the time derivatives in Equation (17) are approximated by and then the defining feature of the 1-SBEM numerical scheme in [10] is how the reaction terms *f, g* in Equation (17) are approximated. The approximations of the reaction terms are given by

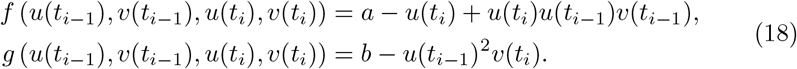

This method is particularly clever as it separates the two unknowns which the FEM solves for in each time step, namely the concentration profiles of the states at the current time step given by *u*(*t*_*i*_) and *v*(*t*_*i*_). This means that in the resulting FEM-formulation, there will only be terms containing inner products between the approximation of the activator *u* and its corresponding test function *ϕ*_1_ as well as inner products between the approximation of the inhibitor *v* and its corresponding test function *ϕ*_2_. Therefore, there is no mixing between the two states and essentially this means that two separate matrix equations can be solved in each time step for both of the states simultaneously which increases the efficiency of the algorithm.

Here, it is important to emphasise that higher values of *γ*, require smaller step sizes *k* in the 1-SBEM-time-stepping procedure. By trial and error, we determined an appropriate step size giving rise to stable solutions for a given eigenmode *n* (Table 1) in the case when *γ* was set to its critical value according to *γ* = *γ*_*c*_(*n*) (16).

**Table 1.** The step sizes *k* used in the 1-SBEM-time-stepping procedure as a function of the eigenmode *n*.

| *Eigenmode, $n$* | $\gamma = \gamma_c(n)$ | *Step size, $k$* |
| --- | --- | --- |
| 1 | 6.87 | 0.01 |
| 2 | 20.62 | 0.01 |
| 3 | 41.24 | 0.005 |
| 4 | 68.73 | 0.005 |

### 4.5 Numerical solutions

For each numerical solution we used initial conditions that were generated by randomly perturbing the steady states concentrations in (3) at every nodal point of the mesh by a value from a uniform distribution on [−10^−4^, 10^−4^). Then, we ran all simulations to a final time of *T* = 50 with single hole located at the South Pole, i.e., at (*x, y, z*) = (0, 0, − 1), with 15 different geodesic hole radii in the range *r* = 0,…, 0.70 where the radius *r* = 0 corresponds to the sphere with no hole. To account for the stochasticity in the initial conditions, we repeated each simulation 20 times, and thus in total we ran

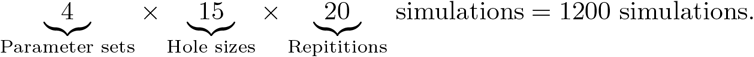

Using all the data that was generated from these simulations, we analysed two types of properties as functions of the increasing hole radius. Firstly, we approximated the final concentration profile of the active component *u* at time *t* = 50 in terms of the first couple of eigenfunctions 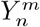 of the Laplace–Beltrami operator (11) in order to deduce what eigenfunctions contributed to the solution. Secondly, we analysed four quantitative metrics of the concentration profile of the active component *u* at time *t* = 50. More specifically, we calculated the number of poles where a pole corresponds to a high–concentration region, the maximum concentration *u*_max_, the area of the pole relative to the total area of the sphere, and the great–circle distance between the midpoint of the introduced hole and the closest pole.

### 4.6 Spectral decomposition of the FEM solution by means of projection

We performed the spectral decomposition of the FEM solution by means of projection onto the spherical harmonics, which form an orthonormal basis for ℒ_2_(*S*^2^). More precisely, let *u*(**x**, *t* = 50) be the FEM solution at time *t* = 50 given by

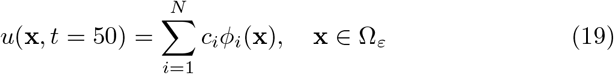

where *N* corresponds to the number of nodes in the mesh, *i* = 1, …, *N* is an index, *c*_*i*_ are coefficients and *ϕ*_*i*_ are the piecewise linear basis functions. The idea is to express the FEM-solution above in terms of the spherical harmonics 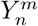 according to

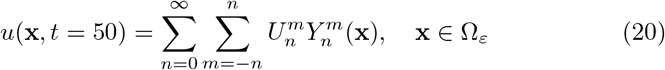

where 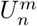 are the unknown coefficients which we wish to compute. We compute the coefficients 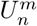 by the following formula

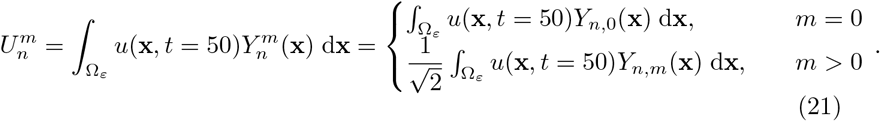

In accordance with [9], we have used the real part of the complex spherical harmonics and taken 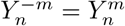. The *Y*_*n,m*_’s in (21) are thus the real spherical harmonics. A table of the *Y*_*n,m*_’s for the eigenmodes *n* = 0, 1, 2, 3, 4, 5 can be found in the Supplementary information (Table S1). For brevity, we only calculate the coefficients 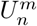 of our FEM-solutions in terms of the first couple of eigenmodes *n*.

### 4.7 Quantifying the number of poles using density based spatial clustering

To calculate the number of poles in an automated fashion, we used the Python based package for machine learning called *scikit-learn* [18]. More precisely, we used the function *DBSCAN*, which stands for *Density-Based Spatial Clustering*, in the following way.

Firstly, we extracted all spatial points **x** ∈Ω_*ε*_ in the mesh that belonged to a pole in the concentration profile *u*(**x**, *t* = 50) which is a high–concentration region. In turn, a point **x** ∈Ω_*ε*_ was classified as belonging to a pole if the concentration at that point in the mesh was greater or equal to 95% of the maximum concentration, i.e., if *u*(**x**, *t* = 50) ≥0.95 *u*_max_. After all coordinates in the mesh that belonged to poles were extracted, the DBSCAN function was implemented in order to find out the number of clusters of spatial points which, in turn, corresponds to the number of poles.

Here, we want to emphasise that the threshold of 95% of the maximum concentration profile is arbitrary. The reason we picked this value was because the number of poles returned by DBSCAN agreed with that obtained by visual inspection.

In addition, the choice of the threshold value to 0.95 *u*_max_ determines the calculated area of the poles relative to the total surface area. This particular threshold value gives rise to a total pole area of approximately 8% of the total surface area. Moreover, a higher threshold value yields a lower total pole area while a lower threshold value yields a higher total pole area.

## Supplementary information

For more results and details, we refer to the Supplementary information associated with this work. We would like to emphasise that the code is available at the public GitHub repository associated with this work [12], and that a large emphasis has been put on writing reproducible Code that is entirely open–source.

## Acknowledgments

JGB would like to thank the Wenner–Gren Foundation for a Research Fellowship and Linacre College, University of Oxford, for a Junior Research Fellowship.

JGB designed and supervised a master thesis [26] which was a prestudy to this work. Here, the effect of holes on the patterns of the Schnakenberg model on a simple planar domain, namely the unit square, was studied by means of FEM simulations.

We would like to thank Professor Christophe Geuzaine from Université de Liège for his help with Gmsh specifically focusing on the Python implementation.

