## Supplementary material for "A hole in Turing’s theory: pattern formation on the sphere with a hole"

### *Supplementary information to the article* A hole in Turing's theory: pattern formation on the sphere with a hole

### S1 FEM-meshes of the unit sphere with a single hole of growing radius

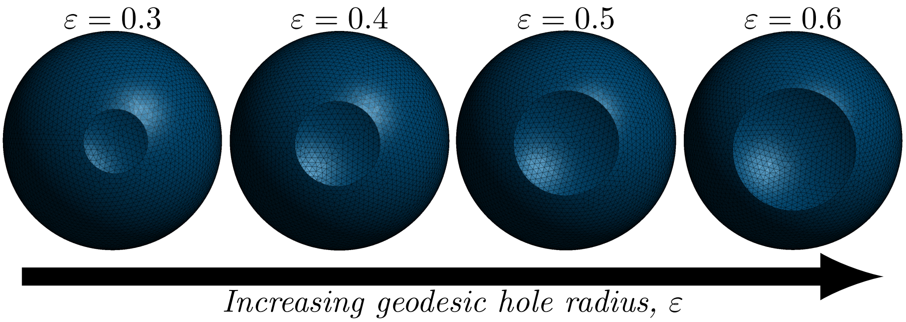


**Fig. S1** *FEM-meshes with an increasing geodesic hole radius $\varepsilon$.* The meshes are presented with a growing area of the holes from left to right where the geodesic radii of the holes are $\varepsilon = 0.3$, $\varepsilon = 0.4$, $\varepsilon = 0.5$ and $\varepsilon = 0.6$, respectively. In our experimental design, we used 15 meshes with a single hole in them corresponding to the geodesic radii $\varepsilon = 0, 0.05, 0.10, \ldots, 0.70$ where $\varepsilon = 0$ corresponds to the mesh without any hole, i.e., the standard unit sphere $S^2$. On all of these meshes, we simulated the Schnakenberg model with stochastic initial conditions, and specifically these initial conditions corresponded to small perturbations around the steady states of the Schnakenberg model. All meshes where generated using *Gmsh* [1] and the FEM simulations were run in *FEniCS* [2, 3].

### S2 The eigenfunctions of the Laplace–Beltrami operator

The eigenfunctions of the Laplace–Beltrami operator on the sphere are the spherical harmonics. They come in various forms, e.g., complex and real, with and without the Condon–Shortley phase factor. Since we set out to replicate the results in [4] to validate our numerical implementation, we chose to use spherical harmonics in accordance to what was used in [4]. This means that we have used the real part of the complex spherical harmonics and taken $Y_n^{-m} = Y_n^m$. The utilized eigenfunctions of the Laplace–Beltrami operator are thus

$$Y_n^m = \begin{cases} Y_{n,0}, & m = 0 \\ \dfrac{1}{\sqrt{2}} Y_{n,m}, & m > 0 \end{cases} \tag{S1}$$

where $Y_{n,m}$ are the real spherical harmonics. We list some of the real spherical harmonics in Table S1 and Table S2. Also, we present a visualisation of the eigenfunctions $Y_n^m$ corresponding to the eigenmodes $n = 0, 1, 2, 3, 4$ when they are projected onto our mesh of the unit sphere (Fig. S2).

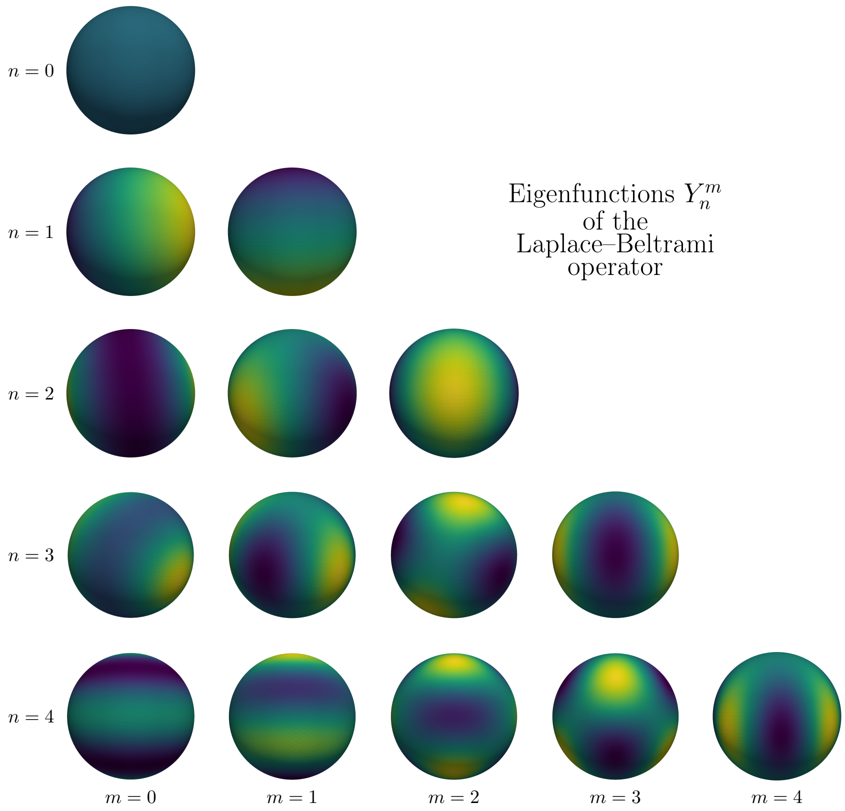


**Fig. S2** *The eigenfunctions $Y_n^m$ of the Laplace–Beltrami operator*. The eigenfunctions $Y_n^m$ corresponding to the eigenmodes $n = 0, 1, 2, 3, 4$ are projected onto the mesh approximating the unit sphere.

**Table S1** The real spherical harmonics corresponding to the eigenmodes $n = 0, 1, 2, 3$.

| *Eigenmode, n* | *Real sph. harm.*, $Y_{n,m}(x, y, z)$, $(x, y, z) \in S^2$, $r = \sqrt{x^2 + y^2 + z^2}$. |
|---|---|
| 0 | $$Y_{0,0}(x, y, z) = \frac{1}{\sqrt{4\pi}} \quad \text{(S2)}$$ |
| 1 | $$Y_{1,0}(x, y, z) = \sqrt{\frac{3}{4\pi}}\frac{z}{r} \quad \text{(S3)}$$ $$Y_{1,1}(x, y, z) = \sqrt{\frac{3}{4\pi}}\frac{x}{r} \quad \text{(S4)}$$ |
| 2 | $$Y_{2,0}(x, y, z) = \sqrt{\frac{5}{16\pi}}\frac{(3z^2 - r^2)}{r^2} \quad \text{(S5)}$$ $$Y_{2,1}(x, y, z) = \sqrt{\frac{15}{4\pi}}\frac{xz}{r^2} \quad \text{(S6)}$$ $$Y_{2,2}(x, y, z) = \sqrt{\frac{15}{16\pi}}\frac{(x^2 - y^2)}{r^2} \quad \text{(S7)}$$ |
| 3 | $$Y_{3,0}(x, y, z) = \sqrt{\frac{7}{16\pi}}\frac{(5z^3 - 3zr^2)}{r^3} \quad \text{(S8)}$$ $$Y_{3,1}(x, y, z) = \sqrt{\frac{21}{32\pi}}\frac{x(5z^2 - r^2)}{r^3} \quad \text{(S9)}$$ $$Y_{3,2}(x, y, z) = \sqrt{\frac{105}{16\pi}}\frac{z(x^2 - y^2)}{r^3} \quad \text{(S10)}$$ $$Y_{3,3}(x, y, z) = \sqrt{\frac{35}{32\pi}}\frac{x(x^2 - 3y^2)}{r^3} \quad \text{(S11)}$$ |

**Table S2** The real spherical harmonics corresponding to the eigenmodes $n = 4, 5$.

| *Eigenmode, n* | *Real sph. harm.*, $Y_{n,m}(x,y,z)$, $(x,y,z) \in S^2$, $r = \sqrt{x^2+y^2+z^2}$. |
|---|---|
| 4 | $$Y_{4,0}(x,y,z) = \sqrt{\frac{9}{256\pi}}\frac{(35z^4 - 30z^2r^2 + 3r^4)}{r^4} \quad (S12)$$ $$Y_{4,1}(x,y,z) = \sqrt{\frac{45}{32\pi}}\frac{x(7z^3 - 3zr^2)}{r^4} \quad (S13)$$ $$Y_{4,2}(x,y,z) = \sqrt{\frac{45}{64\pi}}\frac{(x^2-y^2)(7z^2-r^2)}{r^4} \quad (S14)$$ $$Y_{4,3}(x,y,z) = \sqrt{\frac{315}{32\pi}}\frac{xz(x^2-3y^2)}{r^4} \quad (S15)$$ $$Y_{4,4}(x,y,z) = \sqrt{\frac{315}{256\pi}}\frac{x^2(x^2-3y^2)-y^2(3x^2-y^2)}{r^4} \quad (S16)$$ |
| 5 | $$Y_{5,0}(x,y,z) = \sqrt{\frac{11}{256\pi}}\frac{(63z^5 - 70z^3r^2 + 15zr^4)}{r^5} \quad (S17)$$ $$Y_{5,1}(x,y,z) = \sqrt{\frac{165}{256\pi}}\frac{x(21z^4 - 14z^2r^2 + r^4)}{r^5} \quad (S18)$$ $$Y_{5,2}(x,y,z) = \sqrt{\frac{1155}{64\pi}}\frac{(x^2-y^2)(3z^3-zr^2)}{r^5} \quad (S19)$$ $$Y_{5,3}(x,y,z) = \sqrt{\frac{385}{512\pi}}\frac{(x^3-3xy^2)(9z^2-r^2)}{r^5} \quad (S20)$$ $$Y_{5,4}(x,y,z) = \sqrt{\frac{3465}{256\pi}}\frac{z(x^4-6x^2y^2+y^4)}{r^5} \quad (S21)$$ $$Y_{5,5}(x,y,z) = \sqrt{\frac{693}{512\pi}}\frac{(x^5-10x^3y^2+5xy^4)}{r^5} \quad (S22)$$ |

### S3 The perturbed eigenvalues in the cases $n = 3, 4$

In addition to the results presented in the article, we visualised the perturbed eigenvalues as functions of the hole radius in the cases when $n = 3$ and $n = 4$ (Fig. S3). As can be seen, the parameter choices $d = 18$ and $\gamma = \gamma_c(n)$, where $\gamma_c(n)$ is the critical reaction strength parameter for the eigenmode $n$, trap the perturbed eigenvalues corresponding to the particular value of $n$, when $(a, b) = (0.20, 1.00)$.

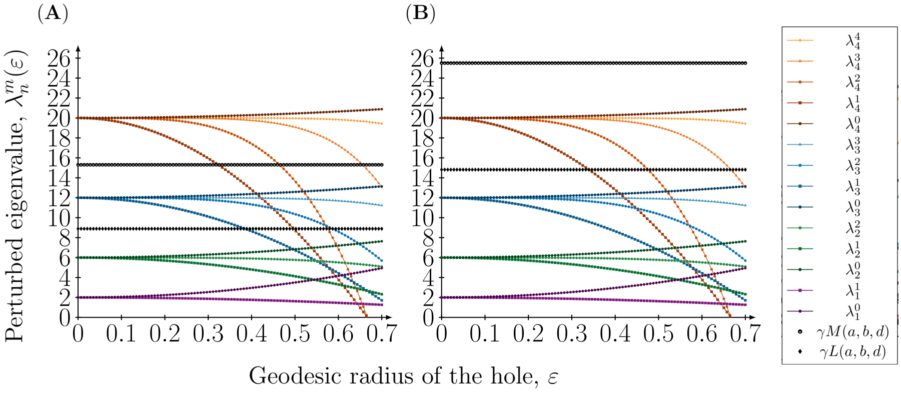


**Fig. S3** *Perturbed eigenvalues as a function of the geodesic radius of the hole on the sphere.* The perturbed eigenvalues $\lambda_n^m(\varepsilon)$ are plotted against the hole radius $\varepsilon$ when $n = 1, 2, 3, 4$ and $m = 0, 1, \ldots, n$. Also, the upper boundary $\gamma M(a, b, d)$ and the lower boundary $\gamma L(a, b, d)$ in the Turing condition involving the eigenvalues giving rise to patterns are illustrated in the dashed lines. The parameters defining these boundaries are chosen to $(a, b) = (0.20, 1.00)$ and the value of $\gamma$ is set to the critical value, i.e., $\gamma = \gamma_c(n)$ for a particular eigenmode $n$. The upper boundary $\gamma M(a, b, d)$ and the lower boundary $\gamma L(a, b, d)$ are illustrated in two cases: (**A**) $(n, \gamma, d) = (3, 41.24, 18.00)$ and (**B**) $(n, \gamma, d) = (4, 68.73, 18.00)$.

### S4 More results from investigating the effect of holes on pattern formation on the sphere

Here, we present more results from the simulations that were run. We used the following parameters of the Schnakenberg model:

$$(a, b) = (0.20, 1.00), \tag{S23}$$

and $\gamma$ was set to its critical value, i.e., $\gamma = \gamma_c(n)$ where $n = 1, 2, 3, 4$ is the particular eigenmode of interest. In this section, we present three types of plots for each parameter set:

1. The spectral coefficients of a single numerical solution $u$ at time $t = 50$ computed on the mesh without a hole (for $n = 1, 2, 3, 4$),

2. The spectral coefficients for all numerical solutions $u$ at time $t = 50$ computed on all meshes with a single hole of increasing radius (for $n = 3, 4$),
3. The quantitative properties for all numerical solutions $u$ at time $t = 50$ computed on all meshes with a single hole of increasing radius (for $n = 3, 4$).

For each parameter set, we run the simulations on 15 meshes, and on each mesh we run 20 repetitions to account for the stochasticity in the initial conditions. In total, this means that we run:

$$\underbrace{4}_{\text{Parameter sets}} \times \underbrace{15}_{\text{Meshes}} \times \underbrace{20}_{\text{Repititions}} \text{ simulations} = 1200 \text{ simulations.}$$

When we run the simulations on the meshes with holes with multiple repetitions, we present the 95%, 50% and 5% quantiles of the quantitative measures.

## S4.1 $(n, d) = (1, 20.00)$

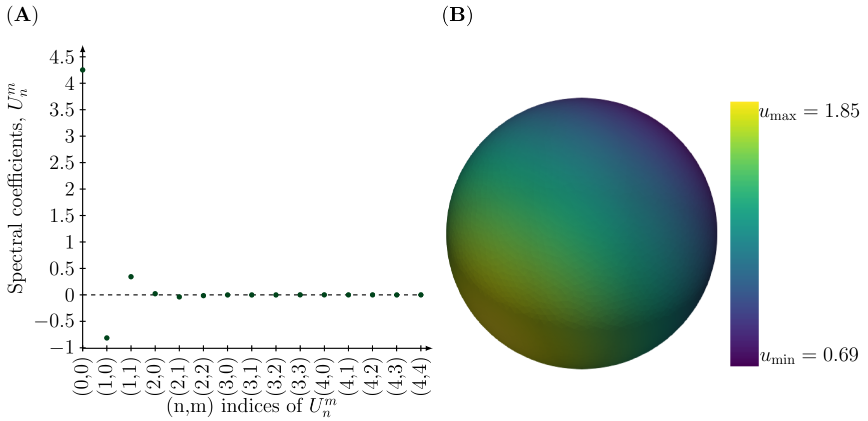


**Fig. S4** *Spectral coefficients of the concentration profile of the active component at time* $t = 50$ *when* $(n, d) = (1, 20)$. The concentration profile $u(\mathbf{x}, t = 50)$, $\mathbf{x} \in S^2$ resulting from the rate parameters $(a, b, d, \gamma) = (0.20, 1.00, 20.00, 6.87)$, where $\gamma = \gamma_c(n = 1)$, is illustrated in two ways. (**A**) The spectral coefficients show that these rate parameters isolate the excited eigenmodes $n = 0$ and $n = 1$. (**B**) The concentration profile of the active component at time $t = 50$ consists of one pole, i.e., high concentration region, and it is bounded between the values $u_{\min} = 0.69$ and $u_{\max} = 1.90$.

## S4.2 $(n, d) = (2, 18.00)$

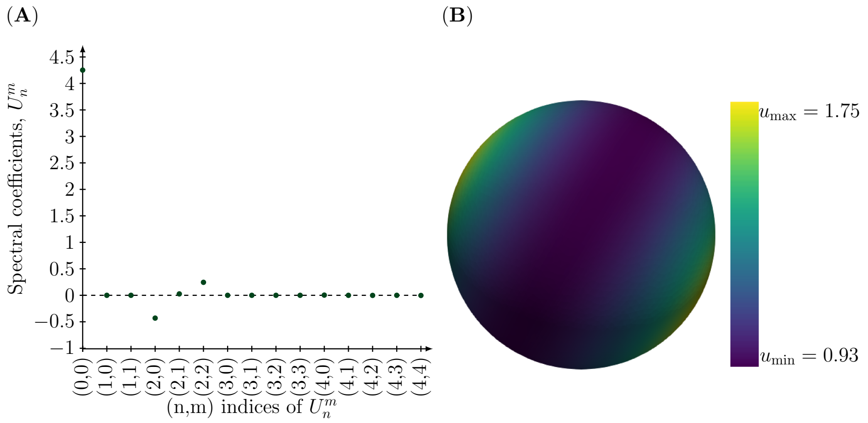


**Fig. S5** *Spectral coefficients of the concentration profile of the active component at time* $t = 50$ *when* $(n, d) = (2, 18)$. The concentration profile $u(\mathbf{x}, t = 50)$, $\mathbf{x} \in S^2$ resulting from the rate parameters $(a, b, d, \gamma) = (0.20, 1.00, 18.00, 20.62)$, where $\gamma = \gamma_c(n = 2)$, is illustrated in two ways. (**A**) The spectral coefficients show that these rate parameters isolate the excited eigenmodes $n = 0$ and $n = 2$. (**B**) The concentration profile of the active component at time $t = 50$ consists of two poles, i.e. high concentration regions, and it is bounded between the values $u_{\min} = 0.93$ and $u_{\max} = 1.70$.

## S4.3 $(n, d) = (3, 18.00)$

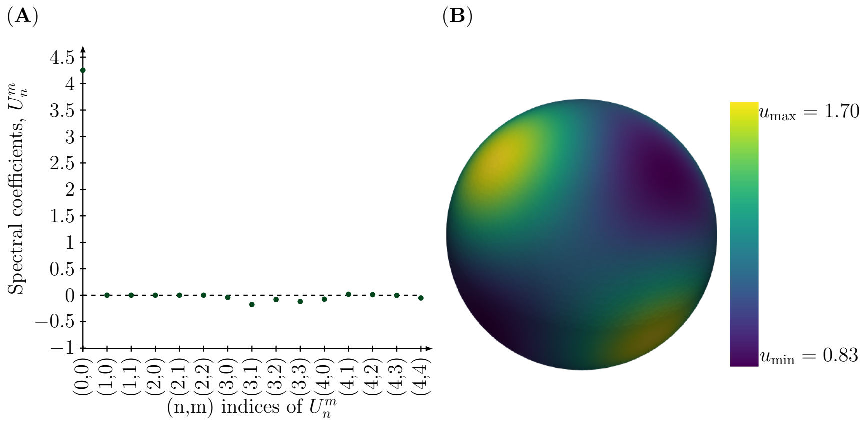


**Fig. S6** *Spectral coefficients of the concentration profile of the active component at time* $t = 50$ *when* $(n, d) = (3, 18)$. The concentration profile $u(\mathbf{x}, t = 50)$, $\mathbf{x} \in S^2$ resulting from the rate parameters $(a, b, d, \gamma) = (0.20, 1.00, 18.00, 41.24)$, where $\gamma = \gamma_c(n = 3)$, is illustrated in two ways. (**A**) The spectral coefficients show that these rate parameters isolate the excited eigenmodes $n = 0$ and $n = 3$. (**B**) The concentration profile of the active component at time $t = 50$ consists of several poles, i.e. high concentration regions, and it is bounded between the values $u_{\min} = 0.83$ and $u_{\max} = 1.70$.

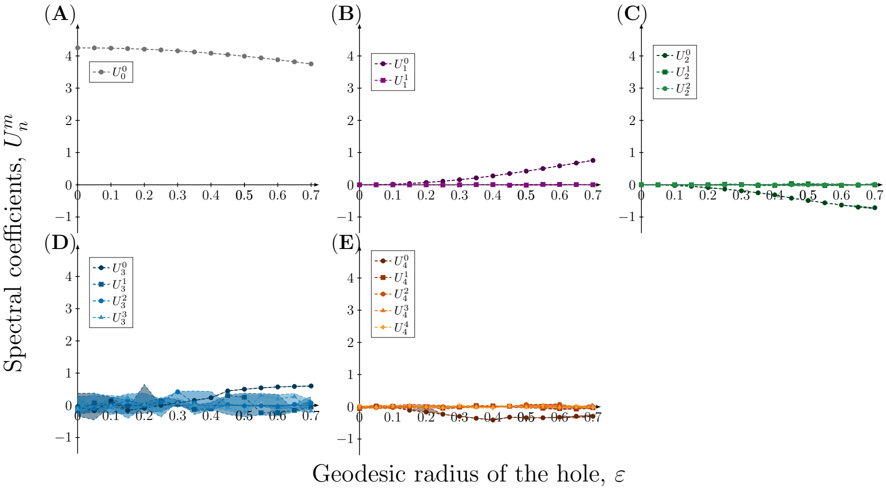


**Fig. S7** *Spectral coefficients of the concentration profile of the active component at time* $t = 50$ *on meshes with a single hole with increasing radius when* $(n, d) = (3, 18)$. The coefficients of the eigenfunctions $Y_n^m$ in the spectral approximation of the concentration profile $u(\mathbf{x}, t = 50),\ \mathbf{x} \in \Omega_\varepsilon$ resulting from the rate parameters $(a, b, d, \gamma) = (0.20, 1.00, 18.00, 41.24)$ are plotted as a function of the geodesic radius of the hole $\varepsilon$ in a few cases. These cases are determined by the coefficients of the eigenfunctions corresponding to the indices: (**A**) $(n, m) = (0, 0)$, (**B**) $(n, m) = (1, 0)$ and $(n, m) = (1, 1)$, (**C**) $(n, m) = (2, 0)$, $(n, m) = (2, 1)$ and $(n, m) = (2, 2)$, (**D**) $(n, m) = (3, 0)$, $(n, m) = (3, 1)$, $(n, m) = (3, 2)$ and $(n, m) = (3, 3)$, (**E**) $(n, m) = (4, 0)$, $(n, m) = (4, 1)$, $(n, m) = (4, 2)$, $(n, m) = (4, 3)$ and $(n, m) = (4, 4)$ and (**F**) $(n, m) = (5, 0)$, $(n, m) = (5, 1)$, $(n, m) = (5, 2)$, $(n, m) = (5, 3)$, $(n, m) = (5, 4)$ and $(n, m) = (5, 5)$. Due to the stochasticity in the initial conditions, each simulation has been repeated 20 times, and to account for the variation in the coefficients the 95%, 50% and 5% percentiles are plotted.

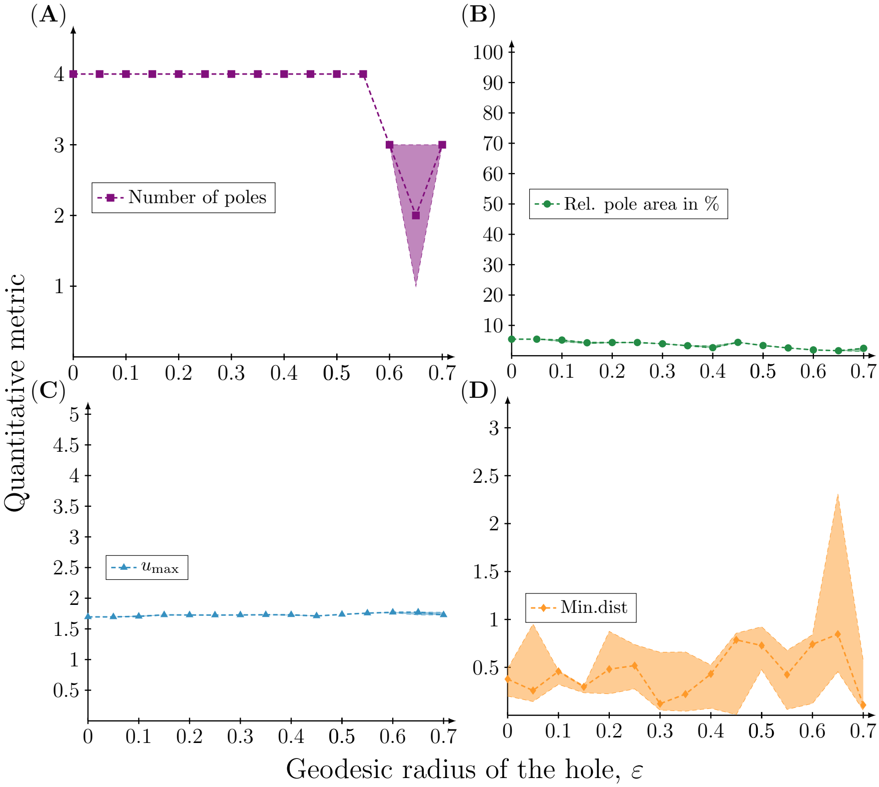


**Fig. S8** *Quantitive metrics of the concentration profile of the active component at time* $t = 50$ *on meshes with a single hole with increasing radius when* $(n, d) = (3, 18)$. Four different quantitative metrics of the concentration profile $u(\mathbf{x}, t = 50)$, $\mathbf{x} \in \Omega_\varepsilon$ resulting from the rate parameters $(a, b, d, \gamma) = (0.20, 1.00, 18.00, 41.24)$ are plotted as a function of the geodesic radius of the hole $\varepsilon$. (**A**) The number of poles corresponding to high concentration regions. (**B**) The total pole area relative to the total surface area. (**C**) The maximum concentration $u_{\max}$. (**D**) The minimal *great circle distance* between a pole and the hole. Each simulation has been repeated 20 times and therefore the 95%, 50% and 5% percentiles of the quantitative metrics are plotted.

## S4.4 $(n, d) = (4, 18.00)$

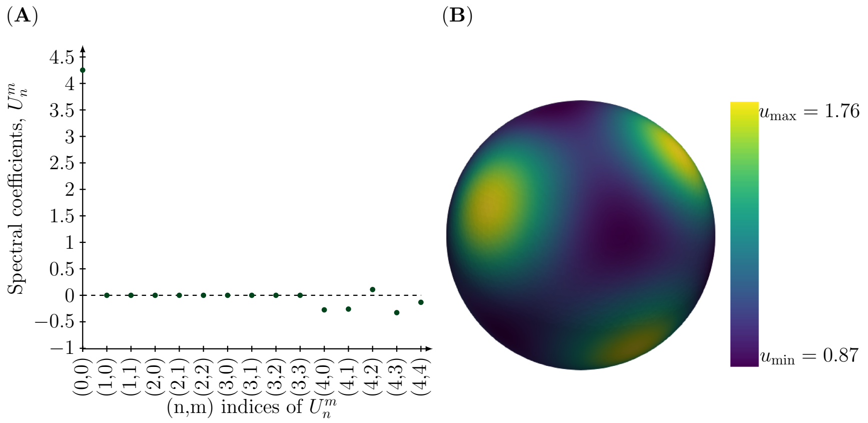


**Fig. S9** *Spectral coefficients of the concentration profile of the active component at time* $t = 50$ *when* $(n, d) = (4, 18)$. The concentration profile $u(\mathbf{x}, t = 50)$, $\mathbf{x} \in S^2$ resulting from the rate parameters $(a, b, d, \gamma) = (0.20, 1.00, 18.00, 68.73)$, where $\gamma = \gamma_c(n = 4)$, is illustrated in two ways. (**A**) The spectral coefficients show that these rate parameters isolate the excited eigenmodes $n = 0$ and $n = 4$. (**B**) The concentration profile of the active component at time $t = 50$ consists of several poles, i.e. high concentration regions, and it is bounded between the values $u_{\min} = 0.87$ and $u_{\max} = 1.80$.

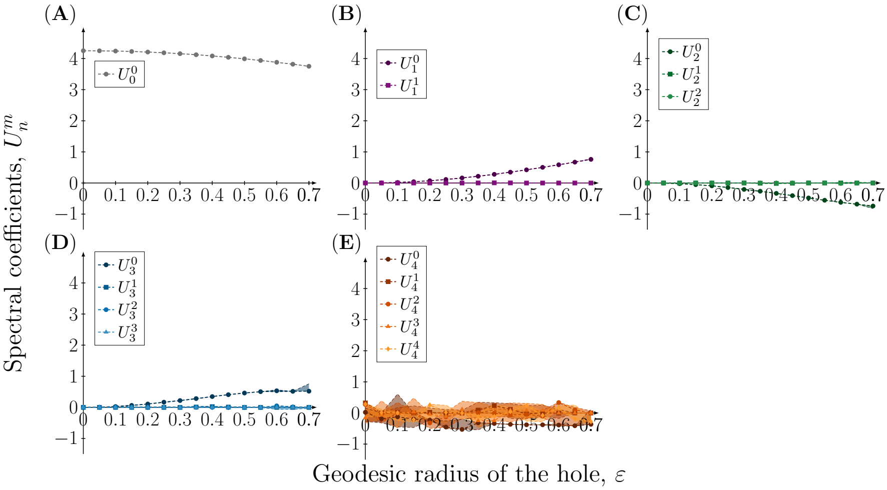


**Fig. S10** *Spectral coefficients of the concentration profile of the active component at time* $t = 50$ *on meshes with a single hole with increasing radius when* $(n, d) = (4, 18)$. The coefficients of the eigenfunctions $Y_n^m$ in the spectral approximation of the concentration profile $u(\mathbf{x}, t = 50),\ \mathbf{x} \in \Omega_\varepsilon$ resulting from the rate parameters $(a, b, d, \gamma) = (0.20, 1.00, 18.00, 68.73)$ are plotted as a function of the geodesic radius of the hole $\varepsilon$ in a few cases. These cases are determined by the coefficients of the eigenfunctions corresponding to the indices: (**A**) $(n, m) = (0, 0)$, (**B**) $(n, m) = (1, 0)$ and $(n, m) = (1, 1)$, (**C**) $(n, m) = (2, 0)$, $(n, m) = (2, 1)$ and $(n, m) = (2, 2)$, (**D**) $(n, m) = (3, 0)$, $(n, m) = (3, 1)$, $(n, m) = (3, 2)$ and $(n, m) = (3, 3)$, (**E**) $(n, m) = (4, 0)$, $(n, m) = (4, 1)$, $(n, m) = (4, 2)$, $(n, m) = (4, 3)$ and $(n, m) = (4, 4)$ and (**F**) $(n, m) = (5, 0)$, $(n, m) = (5, 1)$, $(n, m) = (5, 2)$, $(n, m) = (5, 3)$, $(n, m) = (5, 4)$ and $(n, m) = (5, 5)$. Due to the stochasticity in the initial conditions, each simulation has been repeated 20 times, and to account for the variation in the coefficients the 95%, 50% and 5% percentiles are plotted.

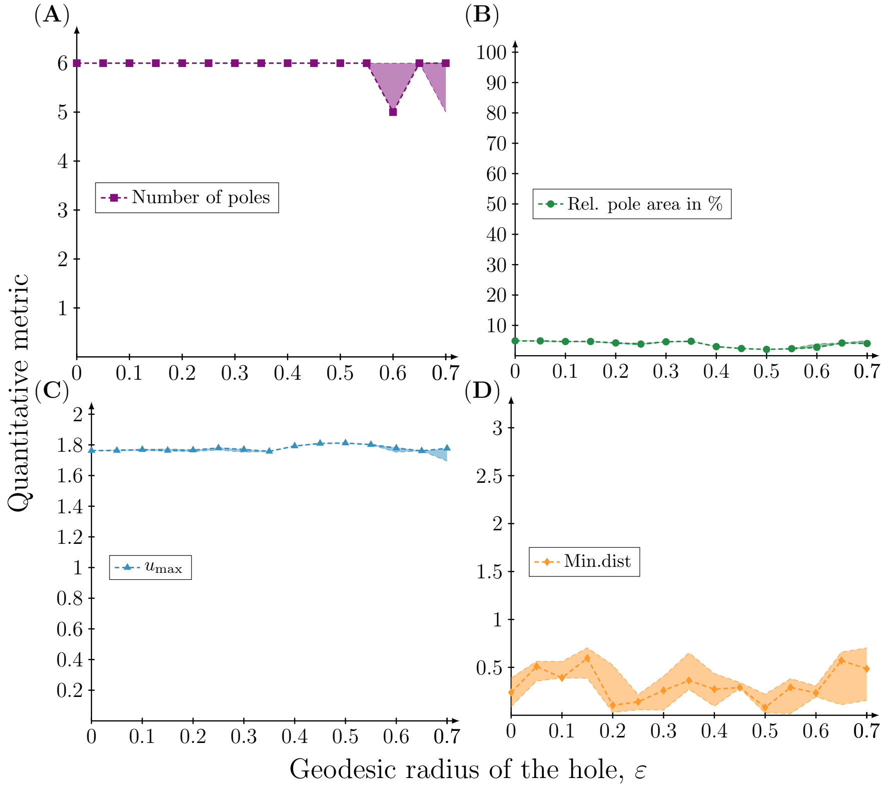


**Fig. S11** *Quantitive metrics of the concentration profile of the active component at time* $t = 50$ *on meshes with a single hole with increasing radius when* $(n, d) = (4, 18)$. Four different quantitative metrics of the concentration profile $u(\mathbf{x}, t = 50)$, $\mathbf{x} \in \Omega_\varepsilon$ resulting from the rate parameters $(a, b, d, \gamma) = (0.20, 1.00, 18.00, 68.73)$ are plotted as a function of the geodesic radius of the hole $\varepsilon$. (**A**) The number of poles corresponding to high concentration regions. (**B**) The total pole area relative to the total surface area. (**C**) The maximum concentration $u_{\max}$. (**D**) The minimal *great circle distance* between a pole and the hole. Each simulation has been repeated 20 times and therefore the 95%, 50% and 5% percentiles of the quantitative metrics are plotted.
